# A discrete sarcomere model reveals internal unbinding dynamics and conditions for regularity of spontaneous oscillatory contractions

**DOI:** 10.1101/2024.07.03.601948

**Authors:** Benjamin Warmington, Jonathan Rossiter, Hermes Bloomfield Gadêlha

**Author notes:** Also at The Soft Robotics Group, Bristol Robotics Laboratory, Bristol (Jonathan Rossiter).

## Abstract

Using a discrete modelling approach for myosin systems we demonstrate how structural differences between single myosin filaments and sarcomeres allow for self similarity during sarcomeric spontaneous oscillatory contractions (SPOC). The form of our modelled SPOC recapitulates the subtleties of *in vitro* SPOCs more closely than prior modelling methods, suggesting we are capturing internal dynamics of the sarcomere that are either not generally considered or previously unknown. These results reinforce the value of discretely modelling molecular motor systems.

---

The half sarcomere is a key element in the hierarchical structure of animal muscle, and is often considered the basic contractile unit (particularly in modelling) [1–4]. It is formed from hundreds of parallel thick myosin cofilaments held in a hexagonal matrix, overlapped with a greater number of passive actin filaments (Fig. 1b) [5, 6]. Sarcomeres are connected in series to form a long filament named a myofibril [5]. When contraction of a muscle is triggered the sarcomeres are flooded with calcium ions (Ca^2+^) [7, 8]. This causes binding sites on actin filaments to be uncovered allowing myosin motors to act on the actin, increasing the overlap and contracting the sarcomeres [8]. In 1978 Fabiato and Fabiato showed that sarcomeres within myofibrils held in isotonic conditions and under partial Ca^2+^ concentration (Fig. 1a) will demonstrate synchronous spontaneous oscillatory contractions (SPOCs) [9]. These spontaneous contractions take the form of a regular sawtooth wave profile, formed from a slow near-linear contraction and rapid extension of the sarcomere (Fig. 1 c) [9–14], and have also been shown to occur at high concentrations of MgADP [12, 15] with isometric and auxotonic boundary conditions [2, 11]. A regular saw tooth has been regularly demonstrated in continuous models of single actomyosin filaments at the limit of a large number of active motors [16–19]. However, the form of the oscillations generated by these models is somewhat distinct from the form of those observed *in vitro* [13, 14], typically demonstrating too non-linear a contraction. This is critical, as the shape of an oscillation is a signature of the non-linearities present in a dynamical system [20], and suggests something is being missed by these models.

**FIG. 1.**
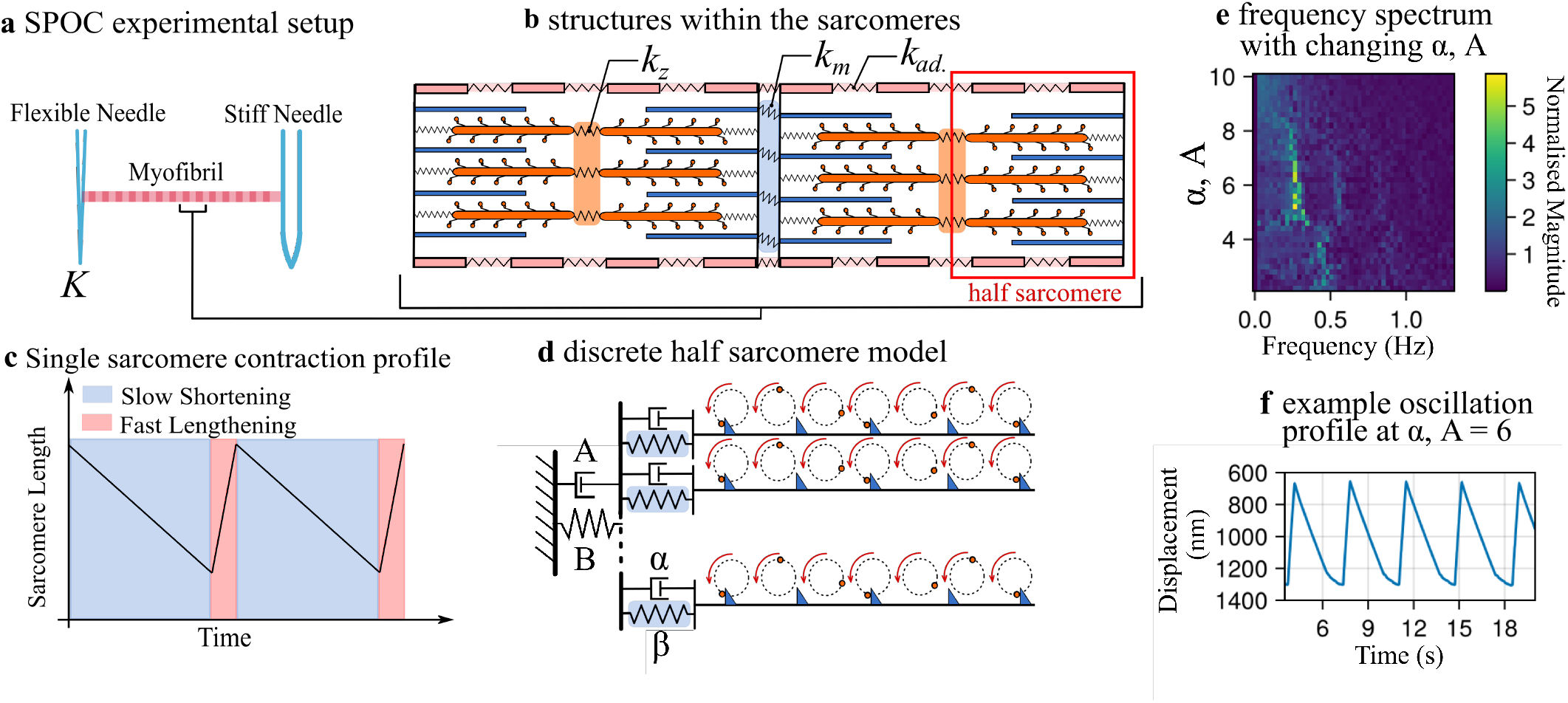
**a** A classic SPOC experiment - a skinned myofibril at partial *Ca*^2+^ activation is held between a stiff needle and a flexible needle [11]. The flexible needle provides an auxotonic boundary. Alternatively, a micro-manipulator can be used at the stiff needle to provide an isotonic boundary [11]. **b** The structure of a sarcomere. Passive thin actin filaments intersect active thick myosin co-filaments. Various auxiliary proteins and structures, as well as the acto-myosin itself provide viscoelastic properties to the sarcomere [5].**c** a typical sarcomere contraction profile, with slow linear/weakly non-linear shortening and fast lengthening [21]. **d** The discrete half sarcomere model. Multiple DCR model filaments [22] are attached in parallel via a spring and dashpot to the ‘Z-line’ by which they are coupled together. This is attached to an additional spring, the auxotonic boundary. **e** Frequency spectrum of model oscillations for range of viscosities, (*α, A*). Other parameters used for the model are; 18 filaments, *B* = 5.4, *β* = 16, N = 56. **f** Example displacement trace of oscillation used to generate **e** at *α* = *A* = 6.

Alongside this no discrete model of myosin or *in vitro* single filament experiment demonstrates regular (self-similar) oscillations [22–24]. These experiments do demonstrate saw tooth oscillations, but with little regularity [23–25]. It is worth noting the rapid unbinding mechanisms for oscillations of ‘minimal actomyosin experiments’ (Refs.[23, 24]) are likely different to those of full sarcomeres [26, 27], however even single actin filaments interacting with a half sarcomere lattice of thick filaments demonstrate highly irregular oscillations [25, 28].

Perhaps one answer to this disparity purely exists in active number of myosin heads, but Warmington *et al*. suggest that even with over 1000 active myosin heads on a single filament regularity is not maintained [22]. Therefore it is likely the more complex mechanical environment auxiliary protein structures [3, 29] and parallelisation of filaments that allows for regularity. Sato *et al* suggests this with a realistic model taking some of the additional mechanical properties into account [26], however, the nature of the model still prevents interrogation of the internal sarcomere dynamics and the role of parallelisation of filaments.

We take a discrete approach to modelling, with parallel independent filaments each containing individually modeled motors. This approach allows for an in-depth consideration of the internal dynamics of the sarcomere that cannot be captured experimentally or are lost in continuous models. Our model shows the requirements for self similar oscillations in a sarcomere-like system, and demonstrates an oscillation shape that captures the subtleties of *in vitro* SPOC more closely than prior modelling methods.

The discontinuously coupled rotor model (DCR model) of myosin demonstrates that many skeletal acto-myosin dynamics can be captured using a minimal rotor model [22]. This abstracts myosin as a rotor which contact equidistantly placed stiffly connected ‘teeth’ representing actin binding sites. Chemical binding is ignored in favour of contact forces between the rotors and teeth. Here we expand upon the DCR by proposing a parallel half sarcomere version of this model, where we investigate additional compliance at the basal end of the actin filaments (Fig. 1 d). Therefore each individual ‘actin filament’ also experiences elastic force at its attachment to an imagined Z plate, simply representing the myriad compliance each separate filament experiences as a single spring per filament. This is the first discretely modelled sarcomere from the myosin level.

We use the same fundamental DCR model as Ref.[22], however instead of using separate forms of the equation for forward and backward movement of the filament (due to saturation of the rotor force upon being backdriven), we adjust the forcing equation of the individual rotors;

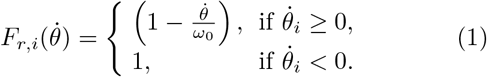

where 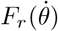 is the rotor’s force in its direction of travel, as a function of its rotational velocity.

The governing equations of the DCR model are given as

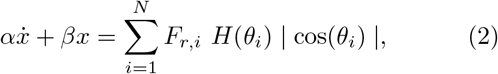

Where *α* is the coefficient of friction, and *β* is a hookian elastic resistance. 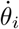 is given by

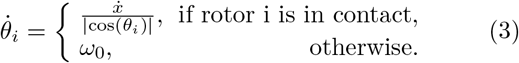

Full descriptions of these equations can be found in Ref.[22].

We take Eq.2 to describe one filament with a ‘compliance’ spring bound to a fixed z plate. We now give the z plate an auxotonic, or springlike, boundary condition, which in *in vitro* experiments is provided by a flexible glass needle or purely the compliance of other sarcomeres [9–14]. We give this boundary the spring constant *B* and the position *X*. We assume the fluid at the imagined Z plate is still viscous if the boundary is a neighboring sarcomere the viscosity could be equal to the internal viscosity. We assign this external viscous resistance ‘*A*’. We can use the spring relations for springs in parallel and series to relate the force on the ‘compliance’ springs to displacement of the boundary. This gives us the overall system of equations for ‘*M* ‘ filaments of ‘*N* ‘ rotors each;

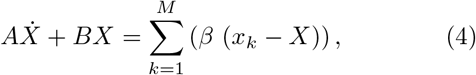

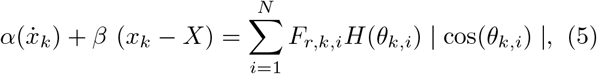

We then solve these equations using a DAE IDA (differential algebraic equation, implicit differential algebraic) solver utilising the Julia differential equations and SUNDIALS packages [30–32]. Code can be found on the authors GITHUB repository.

The DCR model in its original format in Ref.[22] always spontaneously oscillates, whereas a sarcomere requires a reduction in binding affinity in some form to allow SPOC [15]. It is worth noting a myosin motor in *in vivo* conditions may bind to any of the 7 binding sites along a 39nm half helix of actin with reducing affinity as the binding sites turn away from the thick filament [33–35]. In this model binding is limited to one site per half helix, which allows oscillations, and could be considered a reduced affinity. This is arbitrary to a degree, however as binding sites are increased within the DCR model oscillations rapidly die out [22].

We set the number of ‘myosins’ per thick filament, viscous drag and elastic returning force as 56, 6 and 0.4/filament respectively. The number of myosins is an approximation of motor heads available to bind to a thin filament in a half sarcomere (3 directions, approx. 800nm of thick filament in a half sarcomere, 43nm spacing of myosin heads [5, 36–38]). The number of filaments we are able to model in the manner detailed above will be fewer than the actual number of thick and thin filaments within a sarcomere (*>* 100) [5, 36, 39] due to computational constraints. As such the actual elastic returning force found in either the viscoelastic properties of a full sarcomere or Hookian elasticity of a glass needle used in *in vitro* experiments will be too large for this system. To arrive at a ball park estimate, we consider the strength of the auxotonic boundary condition in Ref.[14] (133 *− −*479*nN/µm*) and divide it by the approximate number of thick filaments in the rabbit psoas muscle sarcomere used in Ref.[14] (200*−* 400 [40, 41]). This provides a range of 9.75*·*10^*−*3^*nN/µm* to 73*·*10^*−*3^*nN/µm* or *B* = 0.33 to 2.395 per filament. We set *B* = 0.4 per filament which produces an oscillation of reasonable amplitude. The viscous and protein friction drag within the sarcomere is difficult to set by prior research as it is often contradictory and gives estimates within a large range from the order of *f N· s· m*^*−*1^ [42] to *µN·s ·m*^*−*1^ [18, 23]. As this value is an area of active research, and as of yet unsolved, we set *A* = *α*, and test a range of values to establish which gives the most regular oscillations (Fig. 1e). We find 5 ≤*α* = *A*≥ 7 (≈3.1 *−*4.4*µN· s· m*^*−*1^) gives the most regular oscillations (Fig. 1e). This is on the higher end of our literature range. We use a mid value of this range, *α* = *A* = 6 (Fig. 1f). To gain an overview of large scale dynamics of the model we looked at systems of 1, 2, 3, 6 and 12 filaments at a range of compliance between 2 and 30 (Fig. 2, this figure shows only as low as *β* = 6, but lower values were simulated).

**FIG. 2.**
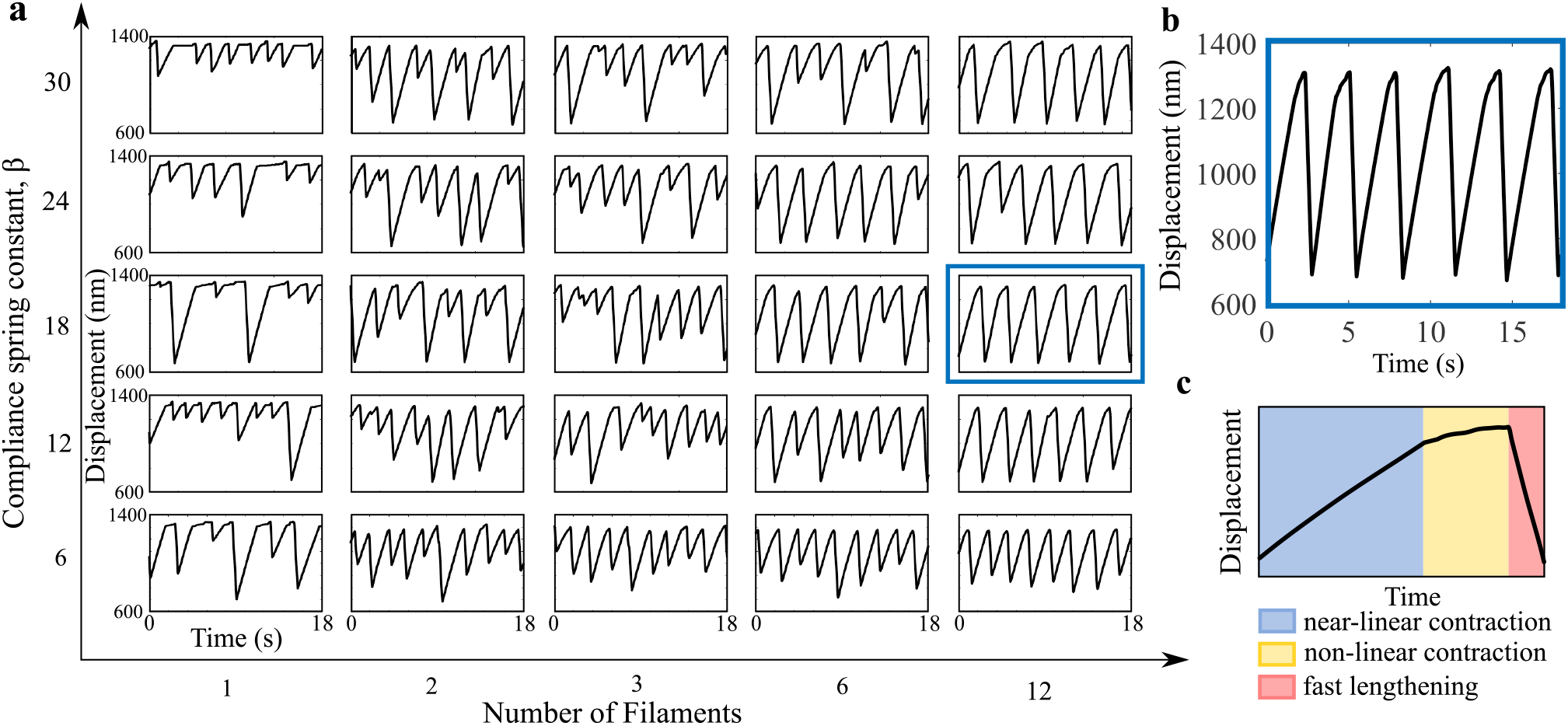
**a** Parameter sweep of discrete half sarcomere, looking at change of behaviour from increasing number of filaments and stiffness of ‘compliance’ spring, *β*. All displacement traces are over 18 seconds and have a range of displacement from 700 1300nm. To ensure the displacement would be approximately the same over increasing numbers of filaments the spring at the auxotonic boundary, ‘B’, is proportional to the number of filaments. **b** Highlighted trace at 12 filaments, with compliance spring constant *β* = 18. This is the most regular of the systems shown. **c** The three sections of an oscillation period. A near linear contraction stage (blue), a non-linear slower contraction stage (yellow) and a fast lengthening stage (red).

Both varying the compliance at the thin filament’s base as well as increasing the number of parallel filaments heavily affects the regularity of the oscillations (Fig. 2a). Qualitatively, all implementations of the model produce typical sawtooth oscillations seen in sarcomeres, however high filament number, medium compliance systems demonstrate the most robustly regular sawtooth with consistent peak, trough and wavelength. All the oscillations are formed of three sections - a slow linear contraction (it appears linear but is in fact weakly non-linear), a non-linear contraction and a fast extension (Fig. 2c. The addition of a third, non-linear component of the oscillation presents a mismatch with the way SPOCs are typically presented in diagram and model [21, 26, 43]. On inspection of experimental *in vitro* results however, this blunted tip is also present in many sarcomere displacement profiles [11, 13, 14] (though not all). We would like to draw particular attention to Fig. 2B of [13], where the *in vitro* wave form is near identical to the oscillations we demonstrate here (we are measuring the displacement of an imagined Z plate rather than sarcomere length, so our oscillations are relatively upside down). It is worth noting the scale of this non-linear oscillation component is shorter or even absent with much lower viscosity, however, as demonstrated in Fig. 1e regularity decreases alongside decreasing viscosity.

Alongside our expectations, it is clear from Fig. 2a increasing the number of filaments greatly increases the regularity of the non-linear contraction of the sarcomere. A closer examination of these sections demonstrates at the limit of linear contraction smooth dynamics of the oscillation break down and are replaced with a seemingly random set of rougher growing steps (Fig. 2c). This section of the wave occurs at approximately the same displacement for every cycle and parameter set. At this section of the wave the separate extensions of the compliance springs (Fig. 3 ii) shows this is the point at which actin filaments start to rapidly stall, reverse and rebind (seen in Figs.3, ii as a thick noisy band beginning at the non-linear wave section). Reverse refers to a point at which a filament has the maximum number of bound ‘myosin’ which can no longer drive the filament, and the unbinding of large numbers of motors allows the filament to backdrive remaining motors and eventually begin to spring the filament backward. The fast extension of a sarcomere can only occur when all of the filaments reverse at once in a catastrophic unbinding event. Figs.3, iii show the motor fraction (proportion of bound motors) decreases at the limit of linear contraction. The abrupt form of this decrease (particularly visible in Fig. 3a, iii) further indicates many motors unbinding at once; individual filaments releasing. The extensions of the compliance springs shows irregular stalling, releasing and rebinding of filaments as irregular oscillations of growing amplitude during the non-linear sarcomere contraction phase (Figs.3, ii). In systems with more filaments these oscillations are averaged out into a smooth and more predictable increase until complete release (Fig. 3c, ii), however with few filaments the irregularities in the unbinding and binding of a single filament constitute a much greater proportion of the overall behaviour, and as such we see less regular oscillations (Fig. 3a, ii). In the minimum system - a single filament - the non-linear section of the oscillation seems to be completely random.

**FIG. 3.**
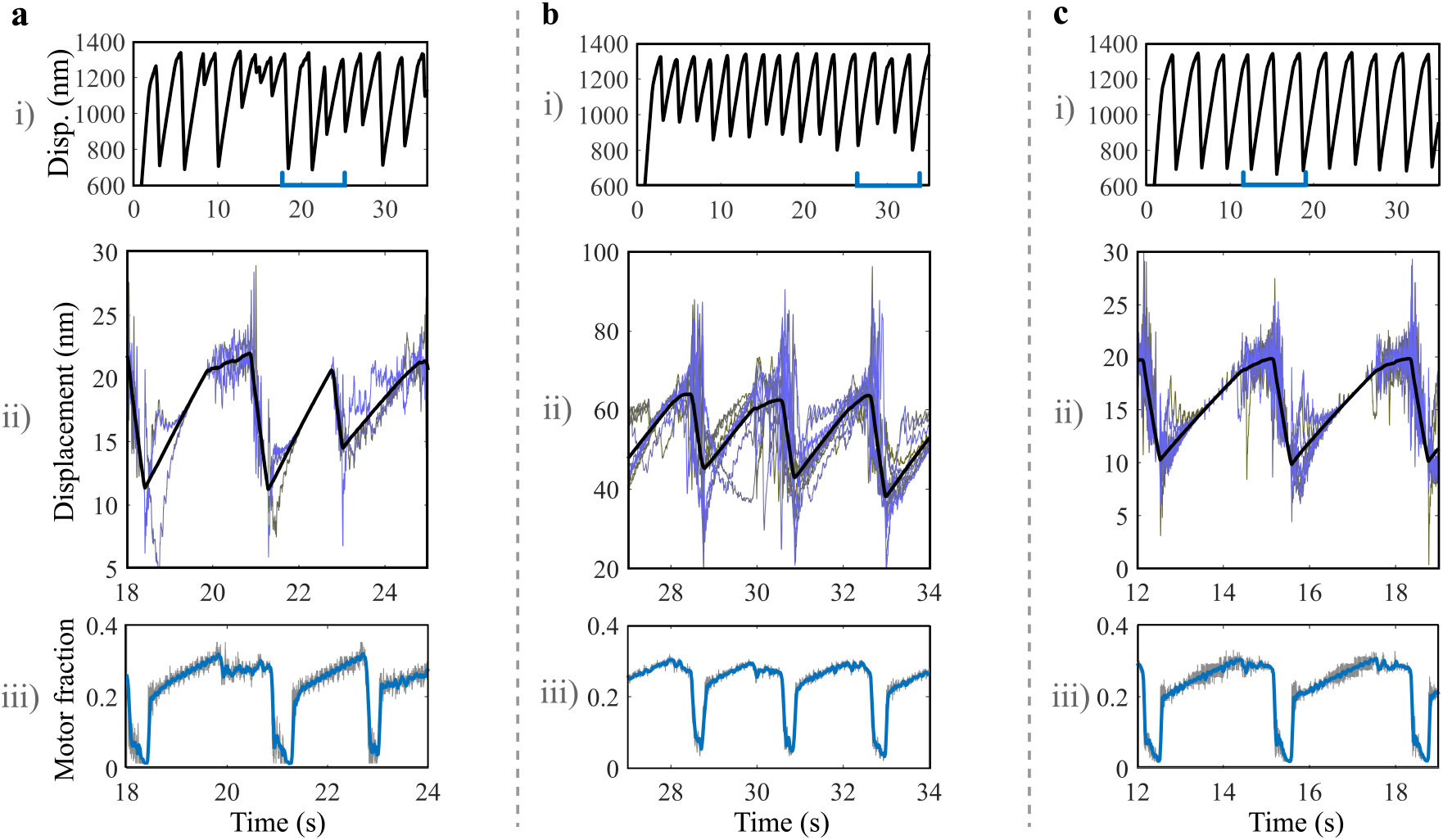
(i) Displacement, (ii) individual ‘compliance’ spring extensions in shades of blue with average in black and (iii) motor fractions for three discrete half sarcomeres. **a** *N*_*f*_ = 3, *β* = 18. The overall displacement is highly irregular, the individual spring extensions show that despite quickly converging to the average displacement there are too few filaments at the cusp of the oscillations to make the cusp regular, with one very protracted cusp and one very short cusp shown here. **b** *N*_*f*_ = 12, *β* = 6 Though these are somewhat regular oscillations the amplitude of the oscillations varies significantly. Individual extensions do not converge to the average at all. The motor fraction follows a fairly regular path, but does not fall as close to zero as stiffer sarcomeres, perhaps contributing to the smaller and more irregular amplitude of the oscillations. **c** *N*_*f*_ = 12, *β* = 18 The most regularly oscillation sarcomere, individual spring extensions quickly converge to the average on each oscillation meaning a similar mechanical environment as the oscillation reaches the cusp of contraction. There are enough filaments contributing to the behaviour at the cusp to help ensure the average remains regular. It is useful to note using the motor fraction that at the limit of linear contraction the number of bound motors decreases, despite the displacement continuing to increase. This is indicative of individual filaments unbinding.

Varying the *β* from low to high suggests neither very compliant nor very stiff architectures of sarcomere can produce regular oscillations. Taking the 12 filament sarcomere explored in Fig. 2, regular oscillations occur only near the middle of the explored values for compliance, with the only highly regular example pictured at *β* = 18. In very compliant systems we see only slightly irregular peaks, with a low amplitude (the lowest value of *β* pictured in Fig. 2 has a peak to peak amplitude of 400nm, however when *β <* 2 it is as small as 150nm) with a short smooth cusp. Very stiff systems demonstrate a greater amplitude of 600nm peak to peak, with a regular limit of linear contraction and rebinding points (Fig. 2, *β* = 30). The irregularity here arises in frequent multi-peaked oscillations, where the rebinding occurs very early, and in the scale of non-linear contraction which varies cusp to cusp from extremely short and pointed to very long and flattened. Intermediate compliance systems also have an amplitude of approx. 600nm peak to peak, however there is a very low occurrence of multi-peaks and, though not perfectly regular, cusps are much more uniform in scale (Fig. 2, *β* = 18). Referring to Fig. 3b, ii and Fig. 3c, ii we can see the difference in behaviour between higher and medium compliance systems. During the fast extension phase of a sawtooth peak in the oscillation profile the subfilaments become separated and rebind at very different extensions. In the high compliance (low *β*) system the filament springs never reconverge to the same extension, meaning the conditions in the system vary peak to peak. Contrasting this with the low and medium compliance systems the extension of all the filaments quite quickly reconverge. This produces similar conditions in the system every oscillation by the time the sawtooth is reaching its zenith, the system is set up for regularity. The regularity difference found between the medium and low compliance systems therefore lies in the non-linear contraction section of the oscillation. As the compliance spring becomes stiffer the sarcomere tends toward the behaviour of a single filament system.

Naturally, this model cannot capture the full level of complexity of an *in vitro* sarcomere. Our base rotating unit is an extreme simplification of a myosin head, and therefore does not encode local biochemistry and kinetics. We only have a single thin filament per thick filament and our model remains one dimensional. In our model we are perhaps demonstrating an idealised sarcomere, with perfectly overlapped thick and thin filaments and a consistent surplus of ATP, held at a calcium concentration that engenders SPOC behaviour. However, it is the first model to show how the complex and varied dynamics of a sarcomeric structure arise from many discrete force producing units. Perhaps more than anything else it demonstrates the minimal complexity required to mechanically produce sarcomere-like dynamics, and reinforces the growing understanding that many complex molecular motor behaviours are predominantly determined by the mechanical environment and architecture in which those motors exist.

This work was funded by the DTP EPSRC, and carried out using the computational facilities of the Advanced Computing Research Centre, University of Bristol - http://www.bristol.ac.uk/acrc/. The authors would also like to thank P. Fuchter for the many discussions on numerical solving of non-linear functions.

